# Recurrent Cysteine Enrichment in Diverse Gut Phage Capsids Suggests Gut-Associated Structural Adaptation

**DOI:** 10.64898/2026.07.03.736451

**Authors:** Riley Anderson, Michael P. Wilczek

## Abstract

**Background:** The gut degrades free virions through low pH, bile salts, and proteases, yet the human virome, dominated by bacteriophages, persists. This persistence may be explained by structural adaptations within the major capsid protein (MCP). Many phage MCPs conserve the HK97 fold, allowing convergent features to be detected across populations.

**Methods:** The MCPs from the Gut Phage Database (GPD) (n=8,478) and INPHARED (n=4,905) were predicted using ProtPhage + Phold and clustered using MMseqs2 (GPD=901 vs INPHARED=606). Structural predictions, conservation analysis, and capsomere modeling were used to characterize cysteine environments.

**Results:** Analysis identified cysteine enrichment in GPD MCPs. Phylogenetic mapping was consistent with convergent evolution of high-cysteine MCPs. Within clusters, 59.6% of cysteine positions were ≥90% conserved; across clusters, 57.2% of high-cysteine MCPs shared at least one structurally conserved cysteine. Simulated N=12 capsomeres showed 83% of cysteines are buried (RSA <10%).

**Conclusions:** These findings suggest gut phages may have convergently evolved cysteine-based capsid stabilization, with implications for engineering therapeutic phages.

## Introduction

The gut virome remains a major research focus due to its complex relationship with human health^1^. This virome, dominated by bacteriophages (phages), persists despite the remarkably hostile environmental conditions it inhabits, such as low pH, bile salts and proteases^1–4^. Phages are important not only for their profound impact on the microbiome, but also for their potential as therapeutic agents, particularly as antibiotic resistance continues to spread^5^. One barrier to oral phage therapy is that administered phages must survive gastric acid and proteolysis to reach the intestine, which can result in premature viral inactivation^6^. As a result, engineering phages, either through directed evolution or genetic modification, has grown in popularity^7,8^. However, optimizing for gut stability remains particularly difficult given the lack of information about which structural or molecular features enable phages to withstand the harsh GI environment^9,10^. Thus, understanding adaptations that allow native gut phages to withstand these stresses may aid the rational design of more stable therapeutic candidates.

The structural stability of phages is driven in part by their capsid, the icosahedral containers that protect the genetic material of the phage. The capsid is comprised of capsomeres, which are the hexameric or pentameric faces made of individual protein subunits (protomers) called major capsid proteins^11^. Despite low sequence similarity, the structure of these protomeric capsid proteins is highly conserved within prokaryotic and some eukaryotic viruses, designated the “HK97 fold”, though other capsid architecture is known to exist^12^. The HK97 fold is comprised of recognizable sub-components. These components include the P-domain, the A-domain, the E-loop, and the N-arm. Notably, the A and P domains are connected by the β-hinge, a three-or-more stranded β-sheet that interfaces the A and P domains ^13^. These subunits are positioned similarly within the oligomeric capsomere, with the A-domain comprising the core of the capsomere and the P-domain facing neighboring capsomeres^14^. The conservation of the HK97 structure within phages makes it an ideal candidate for identifying underlying structural adaptations that contribute to capsid stability.

Most capsid structural studies have focused on either small groups or individual phages ^15,16^. This selective approach limits the ability to detect convergent adaptations across phylogenetically diverse populations. Identifying patterns across large phage groups may elucidate conserved stability features that single-phage approaches would otherwise miss. To address this gap, we performed large-scale sequence and structural analyses of major capsid proteins (MCPs) from phages within the Gut Phage Database (GPD)^3^ and compared them against the INPHARED^17^ reference database to identify features associated with gut-adapted phages. We report significant cysteine enrichment in the MCPs of gut phages, with structural localization patterns in the A and P domains suggesting previously undocumented intra-capsomere stability features.

## Methods and Materials

### Data Preparation and MCP Identification

We designed a MCP identification pipeline using multiple bioinformatics tools. First, files were collected from the GPD online database. The proteome file was downloaded and filtered to remove incomplete genomes using the associated metadata; only genomes with Direct Terminal Repeats (DTR) or Inverted Terminal Repeats (ITR) were retained. The INPHARED database file downloaded from the INPHARED GitHub had the FASTA files trimmed to retain only the sequence identifier for compatibility with downstream tools. Post-filtering phage counts were 11,874 for GPD and 5,411 for INPHARED. Prodigal 2.6.3^18^ was used to produce the GPD and INPHARED proteome files, with N values of 1,000,074 and 540,714 respectively.

The MCPs were predicted using ProtPhage^19^ deployed in Google Colaboratory^20^ using ProtT5-XL-UniRef50 embeddings generated for each proteome. Output was filtered to include only MCPs with a confidence score of 0.99 or above (GPD N=10,569; INPHARED N=6,105). Classification accuracy was validated against 67 curated phage structural proteins from UniProt, containing 58 non-MCP proteins and nine MCP proteins, all nine MCPs correctly identified. ProtPhage-predicted MCPs were then validated using Phold^21^, retaining only proteins independently annotated as “major head protein” or “capsid protein”. After deduplication of identical sequences, this two-level filtering resulted in a final count of 8,478 major capsid proteins for GPD and 4,905 for INPHARED.

The MCP sequences were clustered using MMseqs2^22^ at 50% sequence identity and 80% coverage, consistent with thresholds used for phage protein family grouping^23^. Total clusters were 901 for GPD, and 606 for INPHARED. Cluster representatives were designated by MMseqs2 for further analysis. INPHARED representatives include 576 Caudoviricetes, 23 Malgrandaviricetes, 1 Tectiliviricetes, and 2 unknown based on the taxonomy metadata. GPD representatives are primarily unlabeled, with the 20% of labeled phages all hailing from Caudoviricetes.

### Sequence Analysis and Conservation

After the MCPs were identified and clustered, the representative sequences were analyzed for amino acid composition, protein length, instability index, aliphatic index, GRAVY index, flexibility, and overall charge at pH two and pH seven using BioPython^24^. Fold enrichment of all 20 amino acids was calculated as the ratio of GPD to INPHARED median composition to assess whether compositional differences were specific to cysteine. Proteome-wide cysteine content, or the cysteine percentage, was additionally calculated for each genome as the total number of cysteines divided by the total residues across all proteins in that genome, excluding sequences shorter than 30 residues, and restricted to genomes contributing an MCP representative. Each MCP was then normalized by dividing its cysteine percentage by the proteome-wide value for its own genome, and normalized values were compared between datasets using Wilcoxen rank-sum tests. Cysteine conservation within each cluster was assessed using MAFFT 7.505^25^ with automatic mode selection (--auto). Only clusters with two or more members (404 of 901) were included in conservation analysis; singletons were excluded to avoid false 100% conservation rates.

GC content was calculated for each genome as the proportion of G and C bases over total called bases. To test whether cysteine enrichment could be attributed to GC content, each GPD genome was paired with an INPHARED genome of similar GC content. Pairing was performed by optimal assignment (using the SciPy function: scipy.optimize.linear_sum_assignment), which selects the set of pairs with the smallest total GC difference, and pairs differing by more than one percentage point were discarded. This produced 498 pairs with a mean within-pair GC difference of 0.076 percentage points. Representatives were also divided into GC bands of 10 percentage points, and cysteine content was compared between datasets within each band; bands containing fewer than 15 genomes from either dataset were not tested. Matched and stratified comparisons used Wilcoxon rank-sum tests, the paired comparison additionally used a Wilcoxon signed-rank test, and confidence intervals were obtained by percentile bootstrap with 2,000 resamples.

A phylogenetic tree of both sets of cluster representatives was created using ViPTreeGen^26^, which computes pairwise tBLASTx-based proteomic similarity (SG) scores and constructs a BioNJ tree from the resulting distance matrix (1 − SG). Convergent evolution was assessed by comparing the mean pairwise SG score among all GPD representatives against the mean pairwise SG score among the high-cysteine (≥3 cysteines, top quartile) subset.

### Structural Prediction and Alignment

Structural predictions of all 901 GPD cluster representatives from the MMSeqs2 analysis were generated using ESMFold v1^27^ loaded via PyTorch Hub (facebookresearch/esm) in Google Colaboratory with default parameters. Structures were visualized and domain-colored using UCSF ChimeraX 1.11.1^28^.

Structural alignment was performed using USalign MSTA version 2024.11.08^29^ on cluster representatives containing three or more cysteine residues (top quartile of cysteine content, N=290). Structural alignment was used rather than sequence alignment due to low sequence similarity but high structural conservation of the HK97 fold. Cysteine centroids were defined as alignment columns where at least 15 structures contained a cysteine within ±3 columns.

Centroids within 20-columns were merged to prevent reporting multiple centroids from the same structural region. The same analysis was performed on each of the 20 amino acids for comparison.

### Capsomere Prediction and Cysteine Environment

Capsomere predictions were generated for the twelve clusters with the highest membership that contained at least two cysteines conserved at ≥90% in MAFFT alignments using ColabFold^30^ by inserting six copies of the MCP into the provided query box. Models were generated with default parameters. The top-ranked model by pLDDT (prediction confidence score, 0–100) was used for downstream analysis.

Solvent accessibility was calculated using DSSP^31^ on each predicted capsomere. Relative solvent accessibility (RSA) was derived by dividing absolute accessibility by the theoretical maximum for cysteine (140 Å^2^)^32^. Duplicate cysteines from each protomer were averaged to yield a single value per cysteine site. Cysteine pKa values within capsomeres were predicted using PropKa 3.5.1^33^, pKa values exceeding 50, which indicate ceiling artifacts, were excluded.

Local cysteine environment was characterized using BioPython PDB^24^: for each cysteine Sγ atom, all neighboring residues within five Å were identified and classified as hydrophobic (Ala, Val, Leu, Ile, Met, Phe, Trp, Pro), acidic (Asp, Glu), basic (Lys, Arg, His), or polar (all others). Same-chain and other-chain neighbors were tallied separately. Nearest Sγ–Sγ distances were calculated between each cysteine sulfur atom and all other cysteine sulfur atoms in the capsomere, regardless of chain identity. Values were averaged across six protomer copies to yield a single measurement per cysteine site. The same accessibility analyses were repeated for seven control residue types (Ser, Thr, Ala, Val, Leu, Ile, Met), using the Cβ atom as the reference point and residue-specific maximum accessibility values to allow comparison across residue types.

### Statistical Analysis

Group comparisons were performed using the Wilcoxon rank-sum test, selected because the distributions tested (amino acid composition, pairwise similarity scores, hydrophobic neighbor fractions) are bounded at zero and right-skewed. Effect size for the primary enrichment test was measured using rank-biserial correlation. Figures were produced in R 4.5.2^34^ using packages tidyverse^35^ and patchwork^36^.

All code, metadata, and generated files can be found in the supplementary materials. Claude AI Opus 4.5 and 4.6 were used to assist in generating the code within the supplementary materials.

## Results

To determine whether gut phage MCPs differ in amino acid composition from the INPHARED control, we performed fold enrichment tests using Biopython across all 20 standard amino acids. We identified that the amino acid composition between gut and control phages was largely comparable, except for cysteine, which was 2.39x more abundant in GPD MCPs relative to INPHARED (fig. 1a). We next asked whether this enrichment was an artifact of GC content differences between the datasets. GPD genomes were lower in GC than INPHARED genomes (median 42.3% versus 48.3%), confirming the compositional difference, but cysteine enrichment persisted when GC was controlled. Matching representative INPHARED genomes to GPD genomes on GC content yielded 498 pairs with a mean within-pair difference of 0.076 percentage points, among which GPD MCPs retained 2.35x the cysteine content of their matched INPHARED counterparts. GPD cysteine content per protein remained near 0.5% across the full GC range, while INPHARED content declined with increasing GC and fell to zero above 60%, where the majority of reference phage MCPs contained no cysteine at all (fig. 1b).

**Figure 1.**
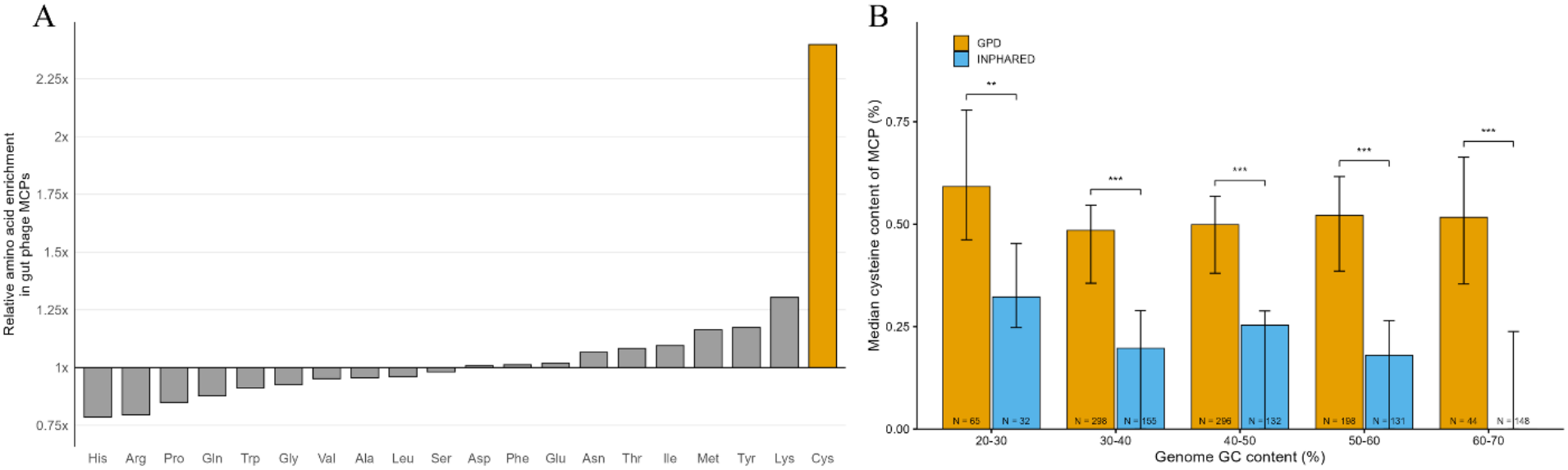
A: Amino Acid Enrichment in Gut Phages Compared to INPHARED Reference Database: Relative enrichment of each amino acid in gut phage major capsid proteins when compared to INPHARED phage major capsid proteins. Bars represent the median amino acid content of GPD phage MCPs divided by the median amino acid content of INPHARED phage MCPs. Below the 1x line indicates the amino acid has greater abundance in INPHARED phage MCPs while above the 1x line indicates greater abundance in GPD phage MCPs. B: Median cysteine content per MCP when compared to genome GC content percentage binned in 10% intervals. Error bars indicate bootstrap confidence intervals. Wilcoxon rank-sum test was used to determine statistical significance; **, p < 0.01; ***, p < 0.001.

To further characterize the cysteine enrichment, we wanted to ensure dominant phage families did not bias the comparison. Thus, we clustered sequences at 50% identity and used one representative per cluster. The 8,478 GPD MCPs and 4,905 reference INPHARED sequences were clustered into 901 and 606 clusters respectively based on sequence similarity.

Representative GPD proteomes carry 1.473x more cysteine than INPHARED genome-wide (fig. 2a), indicating the enrichment is not confined to capsids. MCP specific enrichment between the GPD and INPHARED was 2.39x and was shown to be significant with cysteines making up a median of 0.508% of GPD MCPs and 0.212% of INPHARED (p < 0.0001) (fig. 2b).

**Figure 2.**
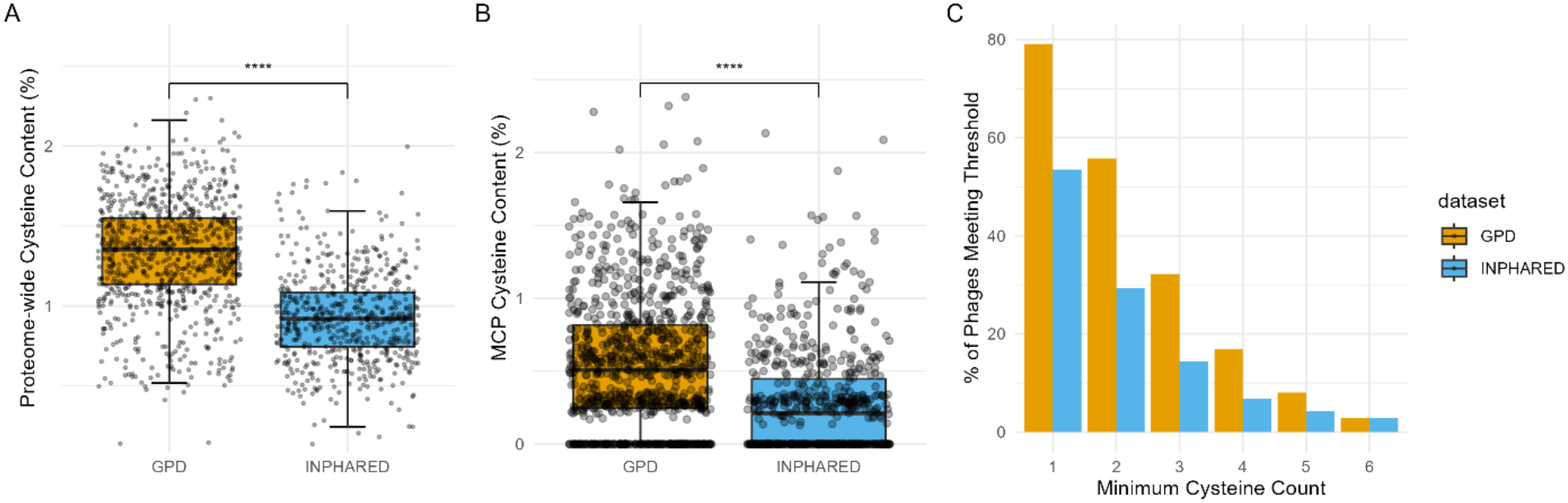
Cysteine Enrichment in Gut Phage Proteomes and Major Capsid Proteins. A: Proteomes from the 901 representative MCP genomes (orange) and 606 reference INPHARED phages (blue). Box and whisker plots represent the distribution of phage proteomes from each database, with the median denoted by the black line and whiskers representing values 1.5 times the distance of the inter-quartile range. B: Percent of major capsid protein comprised of cysteine. Box and whisker plots represent the distribution of phage cysteine content, with the median denoted by the black line and whiskers representing values 1.5 times the distance of the inter-quartile range. Wilcoxon rank-sum test was used to determine statistical significance; ****, p<0.0001. C: Percent of MCPs that contain at least n number of cysteines in 901 gut phage MCPs and 606 reference INPHARED phage MCPs. Fisher exact test was used to determine statistical significance.

Normalizing each MCP to its own genome’s proteome-wide cysteine content leaves a capsid-specific excess of 1.685x (p < 0.0001), showing that capsid cysteine content exceeds what genome composition alone would predict. The MCP enrichment continues across cysteine thresholds, with the percentage of MCPs containing at least one to five cysteines favoring GPD (Fisher’s exact test, all cysteines p < 0.004) until the six-cysteine mark (p = 1.0) (fig. 2c).

To test whether the observed cysteine enrichment better matched with an inherited trait or convergent evolution, GPD and INPHARED cluster representatives were used to construct a phylogenetic tree (fig. 3) based on proteome similarity using VipTreeGen. We compared pairwise proteomic similarity among high-cysteine MCPs (≥3 cysteines) to a size-matched random sample of all GPD representatives. Although this difference reached statistical significance (p = 2.2 × 10^−4^), the mean pairwise similarity values were near-identical (high-cys: 0.00265 vs random: 0.00243), a difference too small to be biologically meaningful. High-cysteine MCPs were therefore no more similar to one another than randomly sampled MCPs are to each other. Visually, gut phage cluster representatives were dispersed throughout the phylogenetic tree rather than forming a single branch, revealing no obvious clustering of high-cysteine MCP phages within a single phylogenetic lineage. Cysteine content was subsequently mapped onto the tree, showing that high-cysteine MCPs overlap primarily though not exclusively with GPD-enriched branches across the phylogeny (fig. 3). A cladogram with labelled tips can be found in the supplementary materials (supplementary Fig. 1).

**Figure 3.**
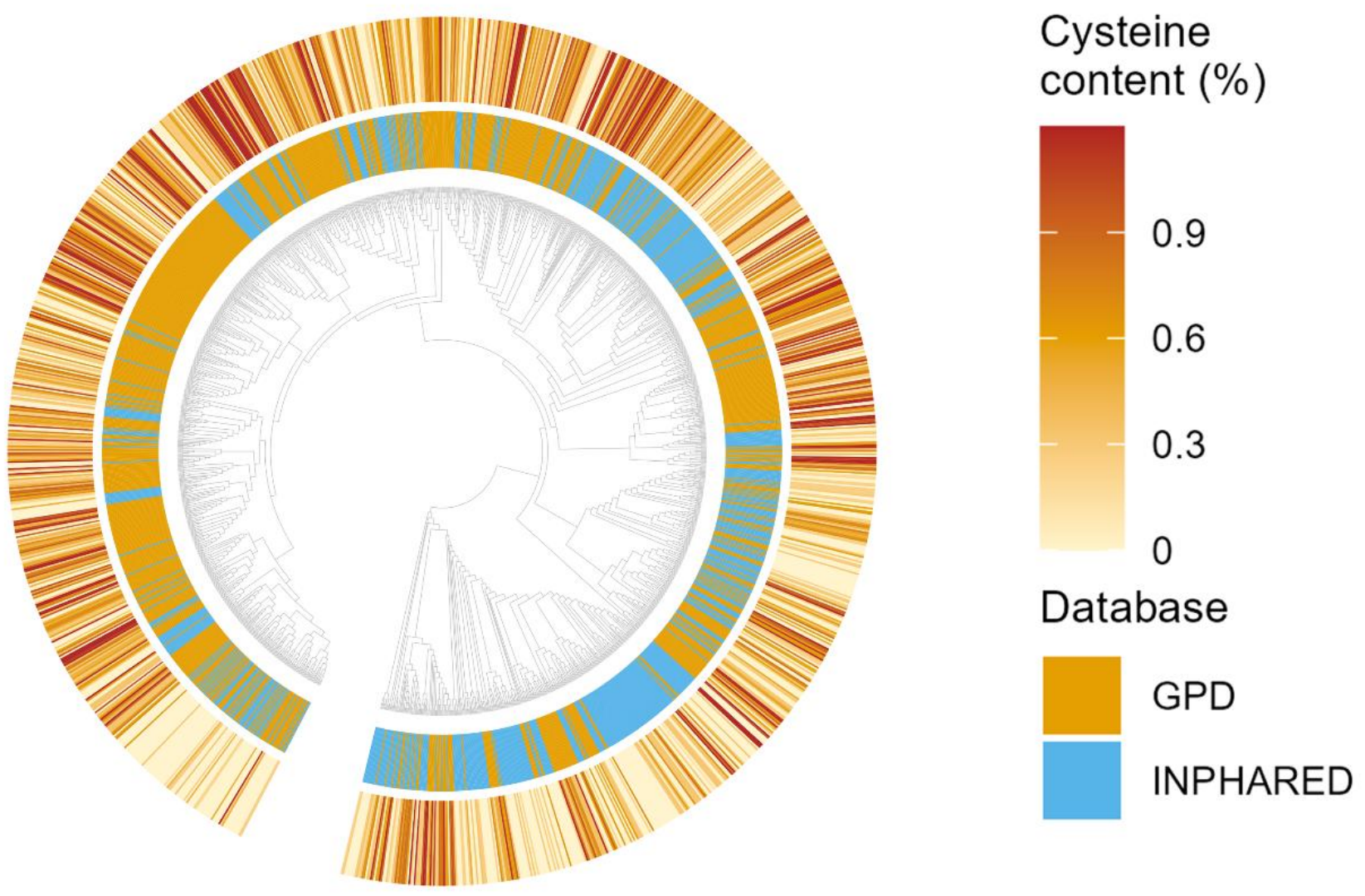
Cysteine Enrichment Mapped onto Cladogram. A circularized cladogram. Each orange (GPD) and blue (INPHARED) line within the inner ring represents a single phage. Phages respective MCP cysteine content by percentage is pictured in the heatmap above it. Heatmap resolves between 0% and 1.2% at maximum.

Having established that high-cysteine phages are phylogenetically dispersed rather than clustered, we next assessed whether cysteine content is randomly accrued or functionally maintained. MAFFT alignment data from each cluster were used to assess cysteine conservation within clusters containing at least two members, comprising 404 of the total 901 clusters. In the largest cluster bin (≥50 members), 38.9% of cysteines were highly conserved, with 16.7% moderately conserved, and 30.6% poorly conserved (fig. 4). This bimodal conservation is consistent with the distribution previously described for cysteine, which is more often either strongly conserved or unconserved than intermediately conserved^37^. Across all 404 clusters and 1,000 cysteine positions, 59.6% of cysteines were highly conserved (≥90% of sequences within a cluster contained a cysteine at the same alignment position), 21.8% were moderately conserved (present in ≥50% of sequences), and 2.7% were poorly conserved (present in <10% of sequences). Though it should be noted that small clusters will skew high in pooled estimates due to the reduced ability to detect variation.

**Figure 4.**
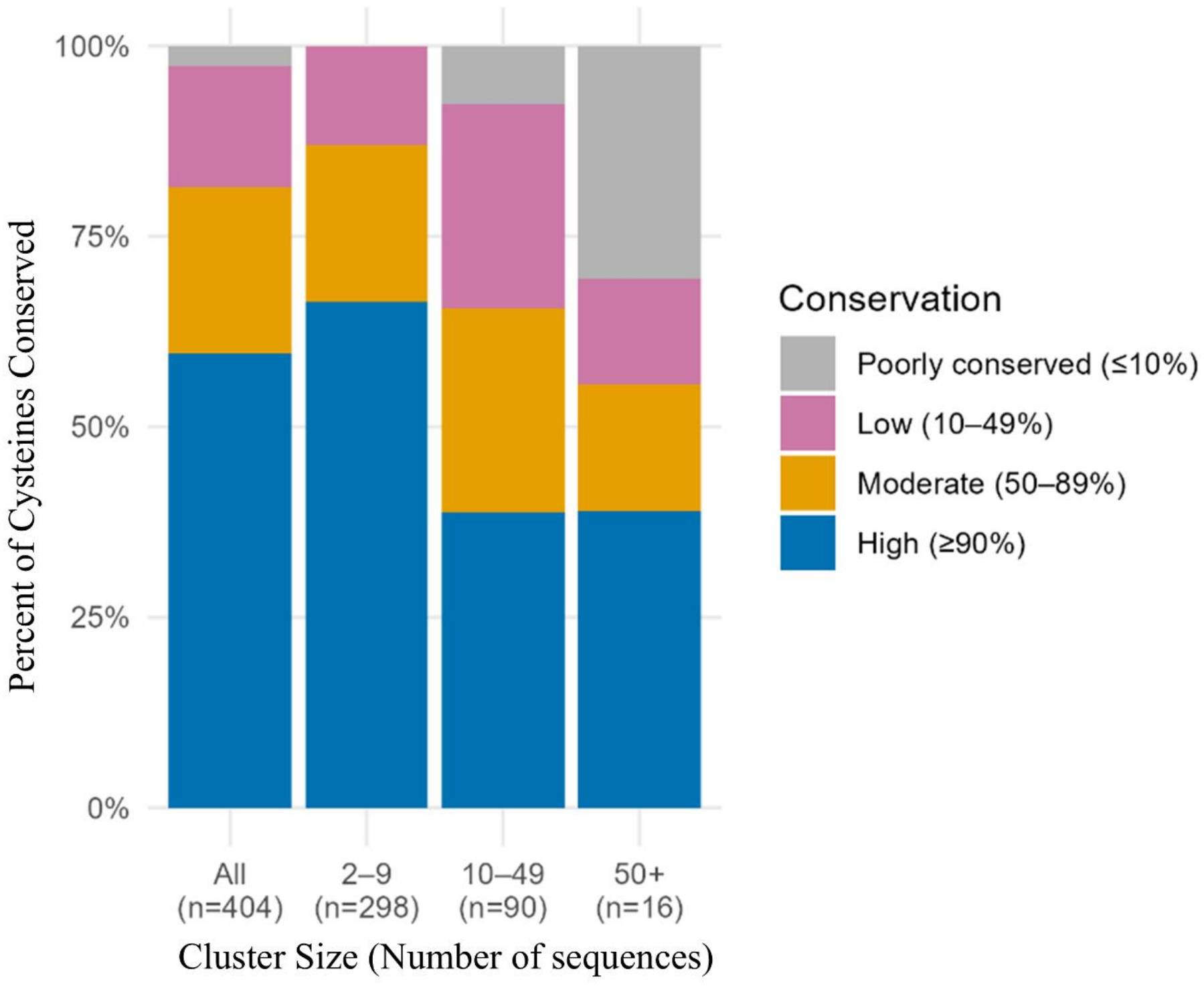
Cysteine Conservation within MCP Clusters. Bar graphs represent conservation of cysteines within clusters of different sizes. N values indicate the number of clusters within each size bin and labels indicate sizes of clusters included in each bin. The “all” bar represents cumulative conservation across all cluster sizes. Blue indicates 90% or higher conservation, orange indicates 50-89% conserved, pink indicates 10-49% conserved, and grey indicates less than 10% conserved.

Next, to assess MCP cysteine conservation between disparate GPD phages, we used USalign structural alignments between the representatives of clusters containing at least three cysteines (top 25% of MCPs) (N= 290). This analysis displayed 12 distinct cysteine centroids (centers of spatial clustering) on GPD gut phage MCPs. Of these centroids, 166 out of 290 high-cysteine MCPs contained a cysteine in at least one of the top three groups (57.2%) (fig. 5A). All centroids had root mean square deviation (RMSDs) below nine Å (fig. 5A), with two having an RMSD below five Å. The majority, 10 of the 12 (83%) of these conserved locations were in the A and P domains (fig. 6), with the remaining two locations landing on the N-terminus arm.

**Figure 5.**
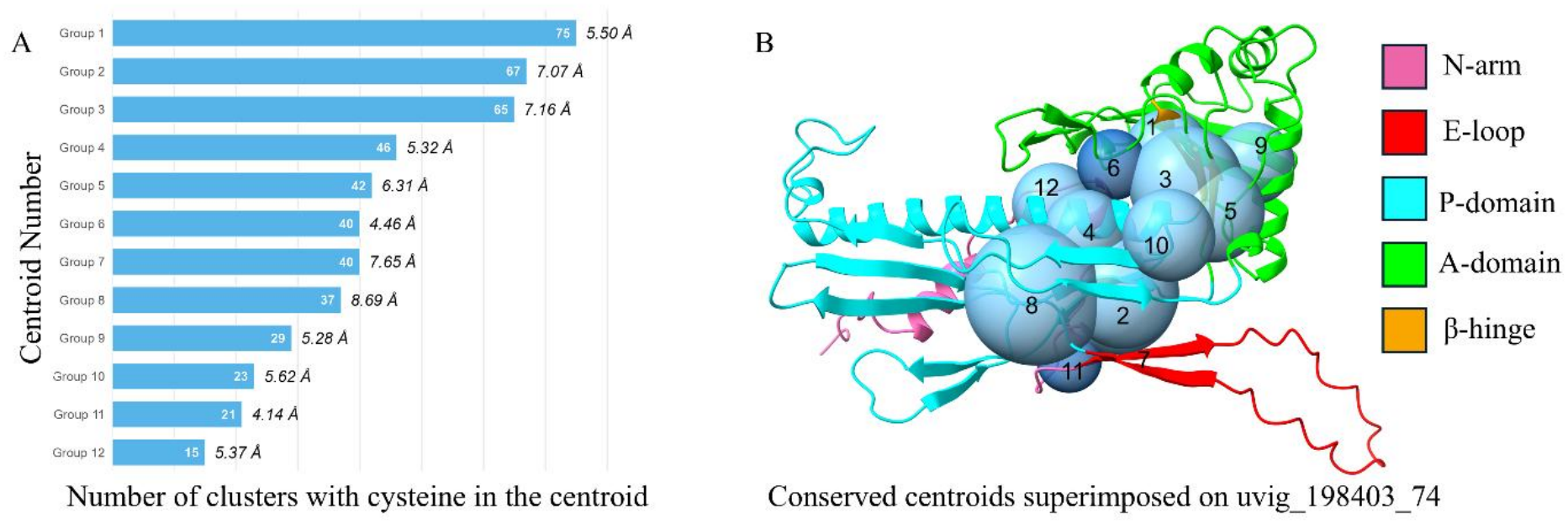
Between Cluster Cysteine Conservation Reveals 12 Distinct Centroids. 290 high cysteine MCP (≥3 cysteines) cluster representatives were structurally aligned with USalign and cysteines within three conservation columns of representative cysteine were considered “conserved” within a centroid. Only centroids with 15 or more cysteines were tracked. A: Number of cluster representatives contributing to a centroid, centroid RMSD in Å is tracked next to the number of representatives. B: Visualization of 12 centroids, labeled 1 to 12, on uvig_198403_74’s HK97 fold domains.

**Figure 6.**
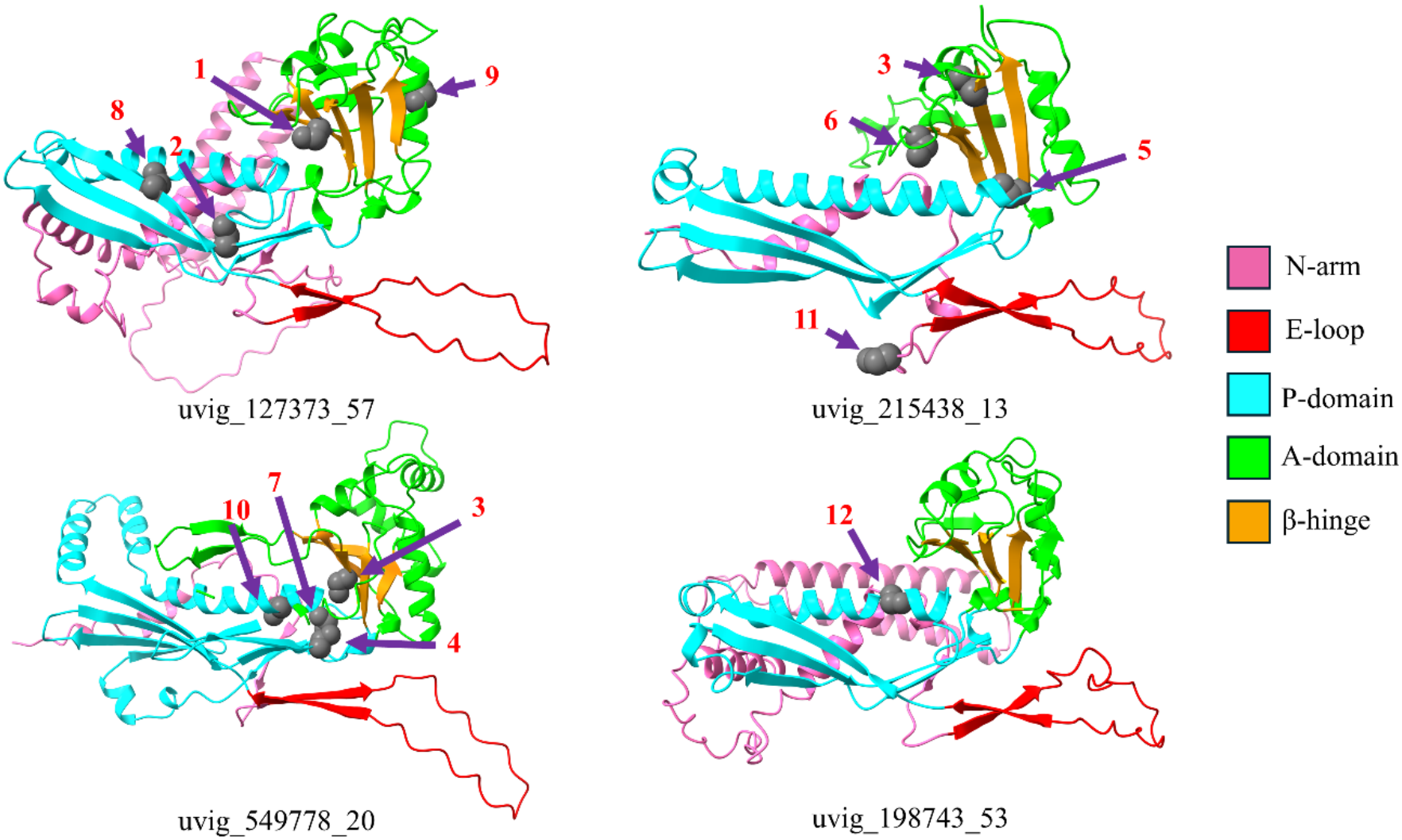
Example Cysteines Contained Within Centroids. Cysteine locations on MCPs that contain a cysteine in each of the centroids one through 12. HK97 fold domains colored to display A and P domain localization pattern. Grey represents a cysteine residue.

Within the A and P domains, four centroids occur on the critical β-hinge region. The same analysis performed on all other 19 amino acids showed that the 6.07% of cysteine residues participating in the top three cysteine centroids was nearly twice the rate of the next-highest amino acid, tryptophan, and well outside the range of the other 19 (mean 2.32%, SD 0.54%) (supplementary fig. 2).

Finally, we were interested in the local protein environment of gut phage MCP cysteines. Because MCPs function in an oligomeric state, representatives from the 12 largest clusters containing at least two highly conserved cysteine residues were used for full capsomere predictions with AlphaFold2-Multimer. Cysteine positions were mapped onto the predicted capsomere assemblies. For each cysteine, relative solvent accessibility (RSA), a measure of surface exposure, predicted pKa, and number of hydrophobic neighbors were calculated (fig. 7). Across all 47 cysteine sites in the 12 predicted structures, 83% were buried (RSA <10%) (fig.7C), 93.5% had a pKa above 9.0, consistent with thiol protonation, and 63.8% were conserved at ≥90% within their cluster. Additionally, cysteines had a median hydrophobic neighbor fraction of 55.6% (fig. 7B), particularly in structured regions with the highest pLDDT values. Comparing seven residues (Ser, Thr, Ala, Val, Leu, Ile, Met) within the same predicted capsomeres, cysteine was the most buried, with a median RSA of 0.77% against 5.65% for leucine, the next most buried, and 27.46% for serine, the closest amino acid analogue (all comparisons p < 1e−4; supplementary fig. 3). This suggests preferential clustering of cysteine residues in highly ordered hydrophobic domains (fig. 7B, C). Only two of the 47 cysteines within predicted capsomeres had any amino acid neighbors from other chains within the capsomere. Most cysteines were isolated; only one thiol out of all 12 structures contained a neighboring sulfur within what is typically associated with disulfide bonding (2.5 Å), with a median distance of 14.65 Å (fig. 7D). Finally, cysteines at β-hinge centroids had 73.0% hydrophobic neighbors compared to 63.1% for non-hinge centroids and 45.9% for non-centroid cysteines (Wilcoxon, W=202, p=0.0037).

**Figure 7.**
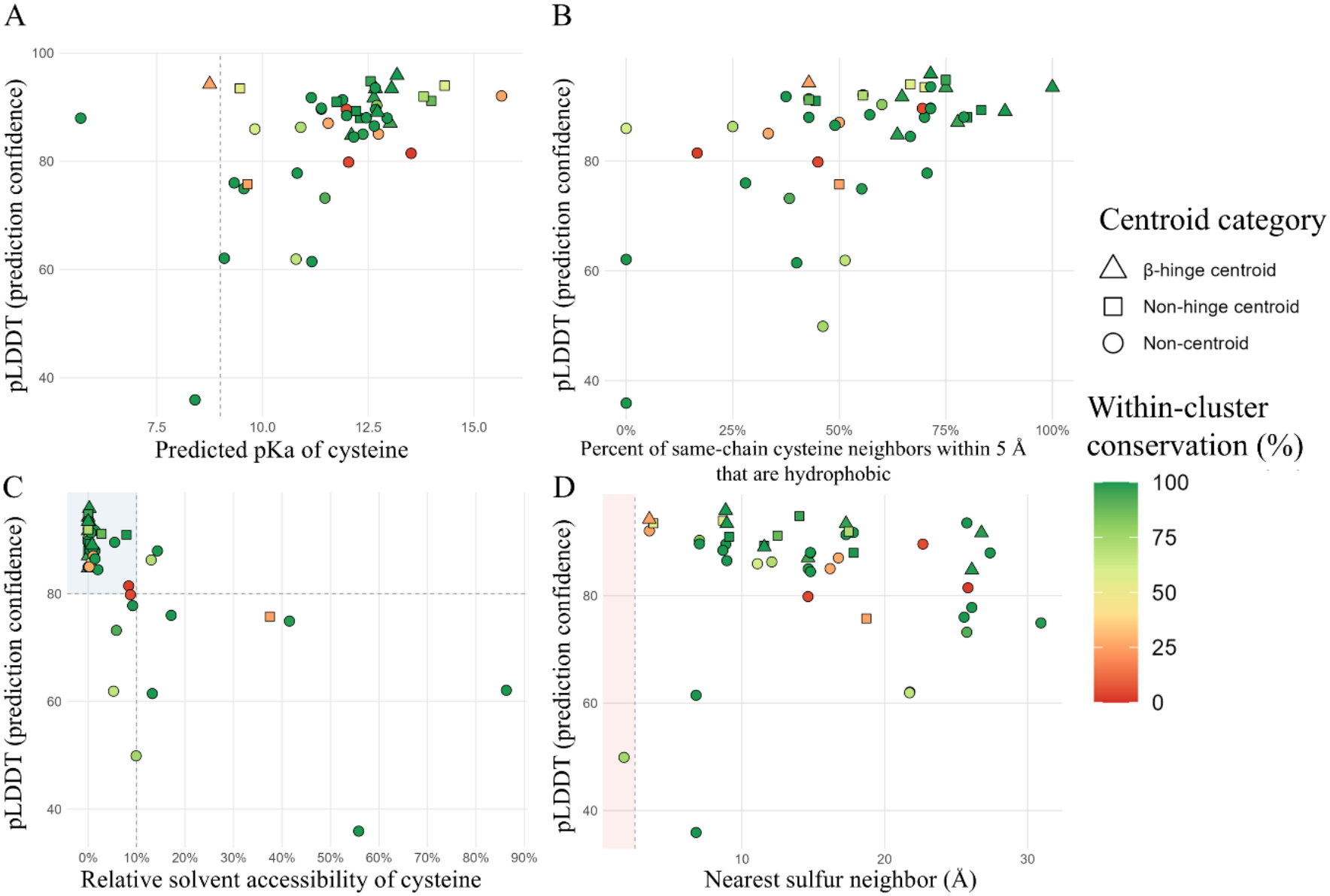
RSA, pKa, and Percent Hydrophobic Neighbors of Cysteines Within Predicted Capsomere Structures. A: Predicted pKa of cysteines within predicted capsomere structures, guideline being the pKa of free cysteine (9.0). B: percent of cysteine neighbors within 5 Å that are hydrophobic. C: Relative solvent accessibility of cysteines within predicted capsomere structures, shaded area represents high-pLDDT cysteines that are buried (≤10% accessible). D: The nearest sulfur neighbor of any given thiol within a capsomere in Å. Red shaded area indicates thiol within 2.5 Å of another sulfur. Color gradient from red to green labels level of conservation within the MCP cluster of origin and the shape indicates centroid status of the cysteine.

## Discussion

To our knowledge, this study represents the first large-scale comparative analysis of a gut phage capsid stability feature, showing that cysteines are significantly enriched in phages within the GPD relative to the INPHARED database (fig. 1, 2). This enrichment is unlikely to be an artifact of GC content, persisting at 2.35x among genomes matched one-to-one on GC and across every GC band tested (fig. 1). Notably, GPD cysteine content remained near 0.5% across the full GC range while INPHARED content declined with increasing GC, a pattern inconsistent with genome composition driving the enrichment. GPD proteomes carry more cysteine genome-wide than INPHARED (1.473x), but capsid enrichment exceeds this baseline, with gut phage capsids sitting 1.685x higher relative to their own proteomes than reference capsids do relative to theirs (fig. 2). High-cysteine phages are phylogenetically dispersed rather than clustered, consistent with recurrent acquisition rather than inheritance from a shared ancestor (fig. 3). The cysteines are generally conserved in both sequence and structure and located primarily in the A and inner P domains (fig. 5, 6). They are also typically buried, protonated, and tend to have amino acid neighborhoods with high hydrophobic content when in their full capsomere context (fig. 7).

While broad enrichment alone suggests non-random placement, the level of conservation found between both related and disparate MCP sequences further indicates that these cysteines are maintained between phage lineages for a structural role. Additionally, the localization of these cysteines to buried, hydrophobic regions of the A and P domains is more consistent with a structural role rather than an exposed functional one. Together, this study suggests that cysteine enrichment may represent a stability adaptation across a diverse range of phages inhabiting the gut.

Traditionally, cysteines are known for their ability to form disulfide bonds. Disulfide bonding within capsids is known to contribute to stability within multiple eukaryotic viruses. HSV-1 virions, for example, whose capsids conserve the HK97 fold, have been shown to require inter-capsomere disulfide bonds to remain stable^38^. Similarly, the major capsid protein of JC polyomavirus, viral protein 1 (VP1), which is prevalent in the human gut during infection^39^, uses transient disulfide bonds to stabilize capsomeres during initial formation and to remain stable once complete^40^. It is worth noting, however, that these eukaryotic viruses acquire disulfide bonds during envelopment within oxidizing compartments, whereas phage capsids assemble inside the reducing environment of the bacterial cytoplasm and are released by lysis. Without interaction with an oxidizing environment, disulfide formation is likely less favorable in phages^41^.

Although disulfide bonding does occur in some phage capsids^42^, cysteines also seem to have alternative roles in protein stability. One proposed role in some globular proteins is hydrophobic packing. While cysteine is traditionally classified as polar, protonated cysteines tend to be generally apolar and are most energetically favorable when contained within hydrophobic pockets^43^. Furthermore, despite its similarity to serine, its closest structural counterpart, isolated cysteines appear to have local environments indistinguishable from hydrophobic amino acids like isoleucine^44^. Recently, the hydrophobicity of free cysteine has also been shown to directly impact intermediate state integrity of the globular protein SPOP^45^, whereas the Rop protein provides evidence of protonated and isolated cysteines reducing the speed of protein unfolding, known as kinetic stability^46^. Perhaps most convincingly, a 2012 study identified a conserved cysteine pair at the AAV capsid pentamer interface buried in a hydrophobic pocket where elevated pKa causes thiol protonation. Substituting one of these residues with valine is tolerated, while substituting serine prevents capsid formation entirely, suggesting the residue’s importance lies in its hydrophobicity rather than disulfide bond formation^47^. Together, these findings provide precedent for hydrophobic packing as a possible primary stability function of cysteine within viral capsids.

Consistent with the reducing environment in which phage capsids assemble, our structural data does not support disulfide bonding as the main stabilizing mechanism in gut phage capsomeres. Firstly, thiol groups were found to be primarily isolated in predicted capsomeres. Only one of the 47 cysteine sites had another sulfur atom within the 2.5 Å typically associated with disulfide formation, and the median distance between sulfur atoms was 14.65 Å, well outside of bonding range (fig. 7D). Notably, the same pipeline placed sulfur atoms from adjacent protomers within 1.8 Å in one capsomere model, indicating that the large distances observed between other thiols do not necessarily reflect a model reluctance to place cysteines in close proximity. Additionally, 83% of tested cysteines were predicted to be primarily or entirely buried, making bonds with external scaffolding proteins unlikely. Even if this were not the case, very few of the cysteines displayed pKa values that would be consistent with deprotonation, even when accounting for the high prediction error value of 4 pKa units associated with PropKa cysteine predictions^48^ (fig. 7A Ultimately, this study proposes that hydrophobic packing at the A–P domain junction may represent an important stabilizing function of the enriched cysteines observed in gut phage MCPs. In predicted capsomeres, cysteines had a median hydrophobic neighbor fraction of 55.6% (fig. 7B), matching the hydrophobic microenvironments previously described for free cysteines in globular proteins and viral capsids^44,47^. Cysteine’s burial exceeded that of every control residue examined, including leucine and its closest structural analogue serine (supplementary fig. 3). The cysteines characterized in this study are also not evenly distributed across the MCP structure. Of the 12 recurring cysteine centroids identified by USalign, ten localize around the junction of the A and P domains (fig. 5, 6). This junction is important for protein folding, particularly the β-hinge, which has been identified as among the most critical structural regions for proper HK97-fold assembly ^13^. Of these ten, centroids one, three, five, and six land directly on the β-hinge, making it the most common identified conservation location across clades, representing 173 clusters or 59.7% of aligned MCPs (fig. 5, 6). The elevated hydrophobic content at β-hinge centroids (73.0% hydrophobic neighbors) relative to non-centroid positions (45.9% hydrophobic neighbors) provides further evidence for this assessment (fig. 7B). The convergence of evidence for cysteine burial, hydrophobic neighborhoods, and preferential localization at the structurally vulnerable A-P junction supports our hypothesis that hydrophobic packing at the β-hinge may help maintain capsomere stability in the harsh gut environment.

Although our work provides a hypothesis supported by multiple lines of convergent evidence, there are several notable limitations. We determined cysteine prevalence and structure within the MCP using multiple bioinformatic analyses and workflow; however, although cysteines are generally conserved in more structured domains, further *in vitro* testing is needed to fully characterize the cysteine microenvironment and its hydrophobicity. Additionally, computational constraints limited capsomere predictions to 12, resulting in a low N value for cysteines in their full oligomeric environment. Future research might focus on predicting capsomeres for all 290 high-cysteine MCPs identified, which would enable more thorough statistical testing on cysteine environments. No *in silico* mutagenesis was performed, so the impact of cysteine loss on structural stability cannot yet be determined; this may be remedied in a future study via molecular dynamics simulations. Finally, no publicly available database of purely non-gut environmental phages exists, making a clean comparison difficult. Indeed, a small proportion of INPHARED representatives (1.2%, 7 of 602) infect obligate gut-associated genera, based on host metadata. Because INPHARED draws from RefSeq, it carries an inherent bias toward clinically relevant phages, such as those infecting Pseudomonas and Mycobacterium.Future work would benefit from a more targeted dataset for environmental control.

In summary, this study finds significant enrichment of cysteine residues in gut phage MCPs from the Gut Phage Database over that of the INPHARED database. These primarily isolated cysteines are predicted to be conserved in hydrophobic regions of the A and P domains, particularly in the β-hinge. While no causal evidence can be provided, multiple lines of correlative evidence suggest that cysteines in these regions engage in hydrophobic packing to stabilize the junction between the A and P domains of protomer MCPs within capsomeres. These findings may inform the rational engineering of phage capsids with enhanced gut stability, an important consideration in the development of therapeutic phages targeting gastrointestinal bacterial infections.

## Supporting information

Supplemental ZIP

## Acknowledgements

Thank you to Dr. Rob Anderson, and Dr. Lucy Anderson for providing manuscript edits. Claude AI (Anthropic, Opus 4.5 and 4.6) was used to assist with code generation and manuscript polish.

Thank you to the Sanger Institute’s GPD database and the INPHARED database for their contribution to this study.

## CITATIONS

1. Cao Z, Sugimura N, Burgermeister E, et al. The gut virome: A new microbiome component in health and disease. eBioMedicine 2022; 81: 104113.

2. Yang I, Nell S, Suerbaum S. Survival in hostile territory: the microbiota of the stomach. FEMS Microbiol Rev 2013; 37: 736–761.

3. Camarillo-Guerrero LF, Almeida A, Rangel-Pineros G, et al. Massive expansion of human gut bacteriophage diversity. Cell 2021; 184: 1098–1109.e9.

4. Almekhlafi SA, Farrag MA, Al-Wahibi MS, et al. Encapsulation of Bacteriophages in Alginate Beads: Improved Viability Under Harsh Simulated Gastric and Intestinal Conditions for Phage Therapy Applications. Pharmaceuticals (Basel) 2026; 19: 363.

5. Lin DM, Koskella B, Lin HC. Phage therapy: An alternative to antibiotics in the age of multi-drug resistance. World J Gastrointest Pharmacol Ther 2017; 8: 162–173.

6. Youssef RA, Sakr MM, Shebl RI, et al. Recent insights on challenges encountered with phage therapy against gastrointestinal-associated infections. Gut Pathog 2025; 17: 60.

7. Meneses L, Valentová L, Santos SB, et al. Directed evolution of phages in biofilms enhances Pseudomonas aeruginosa control through improved lipopolysaccharide recognition. Nat Commun 2025; 16: 10219.

8. Traore K, Seyer D, Mihajlovski A, et al. Engineered bacteriophages for therapeutic and diagnostic applications. Disease Models & Mechanisms 2025; 18: dmm052393.

9. Yin H, Li J, Huang H, et al. Microencapsulated phages show prolonged stability in gastrointestinal environments and high therapeutic efficiency to treat Escherichia coli O157:H7 infection. Vet Res 2021; 52: 118.

10. Xu M, Chen S, Pei H, et al. Engineering bacteriophages for gut health: precision antimicrobials and beyond. J Nanobiotechnology 2026; 24: 62.

11. Gelderblom HR. Structure and Classification of Viruses. In: Baron S (ed) Medical Microbiology. Galveston (TX): University of Texas Medical Branch at Galveston, http://www.ncbi.nlm.nih.gov/books/NBK8174/ (1996, accessed 25 March 2026).

12. Krupovic M, Makarova KS, Koonin EV. Cellular homologs of the double jelly-roll major capsid proteins clarify the origins of an ancient virus kingdom. Proceedings of the National Academy of Sciences 2022; 119: e2120620119.

13. Duda RL, Teschke CM. The amazing HK97 fold: versatile results of modest differences. Current Opinion in Virology 2019; 36: 9–16.

14. Suhanovsky MM, Teschke CM. Nature?s favorite building block: Deciphering folding and capsid assembly of proteins with the HK97-fold. Virology 2015; 479–480: 487–497.

15. Podgorski JM, Freeman K, Gosselin S, et al. A structural dendrogram of the actinobacteriophage major capsid proteins provides important structural insights into the evolution of capsid stability. Structure 2023; 31: 282–294.e5.

16. Stone NP, Demo G, Agnello E, et al. Principles for enhancing virus capsid capacity and stability from a thermophilic virus capsid structure. Nat Commun 2019; 10: 4471.

17. Cook R, Brown N, Redgwell T, et al. INfrastructure for a PHAge REference Database: Identification of Large-Scale Biases in the Current Collection of Cultured Phage Genomes. Phage (New Rochelle) 2021; 2: 214–223.

18. Hyatt D, Chen G-L, LoCascio PF, et al. Prodigal: prokaryotic gene recognition and translation initiation site identification. BMC Bioinformatics 2010; 11: 119.

19. Ou Y, Chen Q, Zhong N, et al. ProtPhage: a deep learning framework for phage viral protein identification and functional annotation. Brief Bioinform 2025; 26: bbaf285.

20. Bisong E. Google Colaboratory. In: Bisong E (ed) Building Machine Learning and Deep Learning Models on Google Cloud Platform: A Comprehensive Guide for Beginners. Berkeley, CA: Apress, pp. 59–64.

21. Bouras G, Grigson SR, Mirdita M, et al. Protein structure-informed bacteriophage genome annotation with Phold. Nucleic Acids Res 2026; 54: gkaf1448.

22. Steinegger M, Söding J. MMseqs2 enables sensitive protein sequence searching for the analysis of massive data sets. Nat Biotechnol 2017; 35: 1026–1028.

23. Gauthier CH, Cresawn SG, Hatfull GF. PhaMMseqs: a new pipeline for constructing phage gene phamilies using MMseqs2. G3 (Bethesda) 2022; 12: jkac233.

24. Cock PJA, Antao T, Chang JT, et al. Biopython: freely available Python tools for computational molecular biology and bioinformatics. Bioinformatics 2009; 25: 1422–1423.

25. Katoh K, Misawa K, Kuma K, et al. MAFFT: a novel method for rapid multiple sequence alignment based on fast Fourier transform. Nucleic Acids Res 2002; 30: 3059–3066.

26. Nishimura Y, Yoshida T, Kuronishi M, et al. ViPTree: the viral proteomic tree server. Bioinformatics 2017; 33: 2379–2380.

27. Lin Z, Akin H, Rao R, et al. Evolutionary-scale prediction of atomic-level protein structure with a language model. Science 2023; 379: 1123–1130.

28. Pettersen EF, Goddard TD, Huang CC, et al. UCSF ChimeraX: Structure visualization for researchers, educators, and developers. Protein Science 2021; 30: 70–82.

29. Zhang C, Shine M, Pyle AM, et al. US-align: universal structure alignments of proteins, nucleic acids, and macromolecular complexes. Nat Methods 2022; 19: 1109–1115.

30. Mirdita M, Schütze K, Moriwaki Y, et al. ColabFold: making protein folding accessible to all. Nat Methods 2022; 19: 679–682.

31. Hekkelman ML, Salmoral DÁ, Perrakis A, et al. DSSP 4: FAIR annotation of protein secondary structure. Protein Sci 2025; 34: e70208.

32. Miller S, Janin J, Lesk AM, et al. Interior and surface of monomeric proteins. Journal of Molecular Biology 1987; 196: 641–656.

33. Olsson MHM, Søndergaard CR, Rostkowski M, et al. PROPKA3: Consistent Treatment of Internal and Surface Residues in Empirical pKa Predictions. J Chem Theory Comput 2011; 7: 525–537.

34. R: The R Project for Statistical Computing, https://www.r-project.org/ (accessed 29 May 2026).

35. Wickham H, Averick M, Bryan J, et al. Welcome to the Tidyverse. Journal of Open Source Software 2019; 4: 1686.

36. Pedersen TL. patchwork: The Composer of Plots, https://cran.r-project.org/web/packages/patchwork/index.html (2025, accessed 29 May 2026).

37. Marino SM, Gladyshev VN. Cysteine function governs its conservation and degeneration and restricts its utilization on protein surface. J Mol Biol 2010; 404: 902–916.

38. Szczepaniak R, Nellissery J, Jadwin JA, et al. Disulfide Bond Formation Contributes to Herpes Simplex Virus Capsid Stability and Retention of Pentons▿. J Virol 2011; 85: 8625–8634.

39. Shadbash P, Hosseini SM, Shoraka S, et al. Possible association between polyomaviruses and gastrointestinal complications: a narrative review. Gastroenterol Hepatol Bed Bench 2024; 17: 121–131.

40. Kobayashi S, Suzuki T, Igarashi M, et al. Cysteine Residues in the Major Capsid Protein, Vp1, of the JC Virus Are Important for Protein Stability and Oligomer Formation. PLoS One 2013; 8: e76668.

41. Pawlowski A, Moilanen AM, Rissanen IA, et al. The Minor Capsid Protein VP11 of Thermophilic Bacteriophage P23-77 Facilitates Virus Assembly by Using Lipid-Protein Interactions. J Virol 2015; 89: 7593–7603.

42. Sonani RR, Esteves NC, Horton AA, et al. Neck and capsid architecture of the robust Agrobacterium phage Milano. Commun Biol 2023; 6: 921.

43. Iyer BR, Mahalakshmi R. Hydrophobic Characteristic Is Energetically Preferred for Cysteine in a Model Membrane Protein. Biophysical Journal 2019; 117: 25–35.

44. Nagano N, Ota M, Nishikawa K. Strong hydrophobic nature of cysteine residues in proteins. FEBS Letters 1999; 458: 69–71.

45. Pagano L, Diop A, Pennacchietti V, et al. A single buried cysteine acts as a hydrophobic stabilizer of a folding intermediate and transition state in the MATH domain of SPOP. Protein Sci 2025; 34: e70138.

46. Hari SB, Byeon C, Lavinder JJ, et al. Cysteine-free rop: A four-helix bundle core mutant has wild-type stability and structure but dramatically different unfolding kinetics. Protein Science 2010; 19: 670–679.

47. Pulicherla N, Kota P, Dokholyan NV, et al. Intra- and Inter-Subunit Disulfide Bond Formation Is Nonessential in Adeno-Associated Viral Capsids. PLOS ONE 2012; 7: e32163.

48. Awoonor-Williams E, Golosov AA, Hornak V. Benchmarking In Silico Tools for Cysteine pKa Prediction. J Chem Inf Model 2023; 63: 2170–2180.

