## Supplementary figures and images for "Recurrent Cysteine Enrichment in Diverse Gut Phage Capsids Suggests Gut-Associated Structural Adaptation"

### Supp. Figure 1.pdf

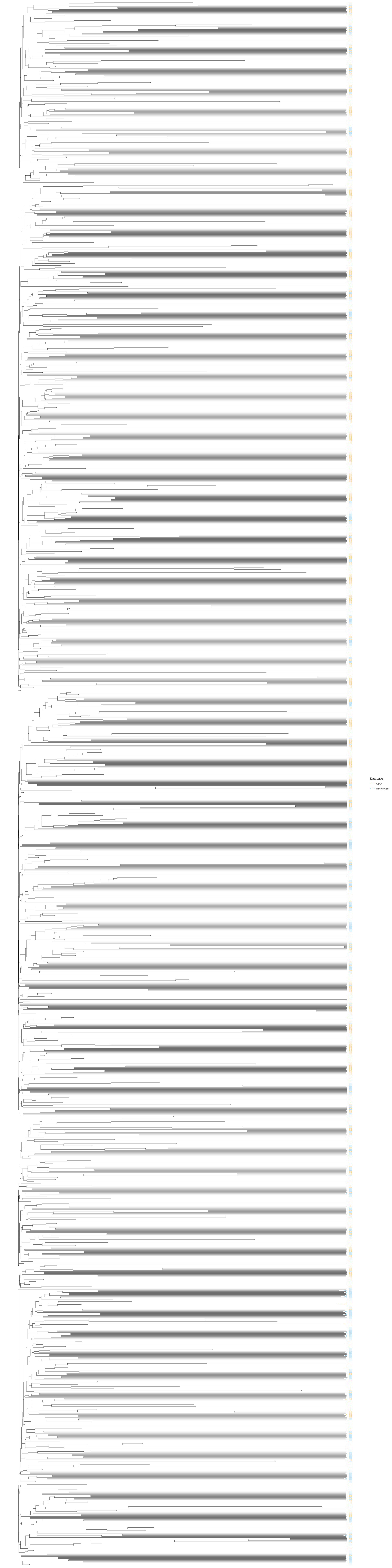
Database
GPD
INPHARED

### Supp. Figure 2.png

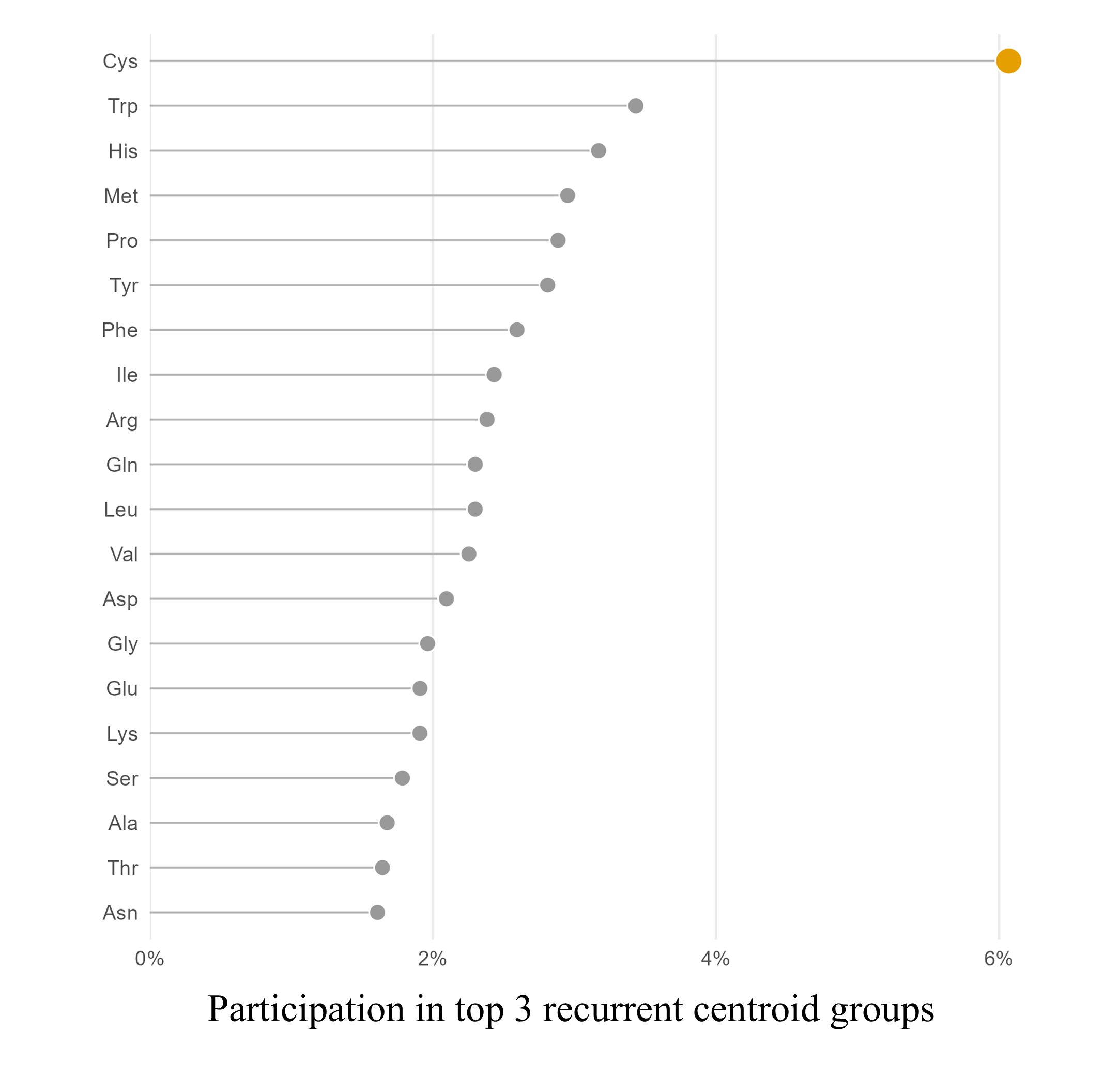

### Supp. Figure 3.png

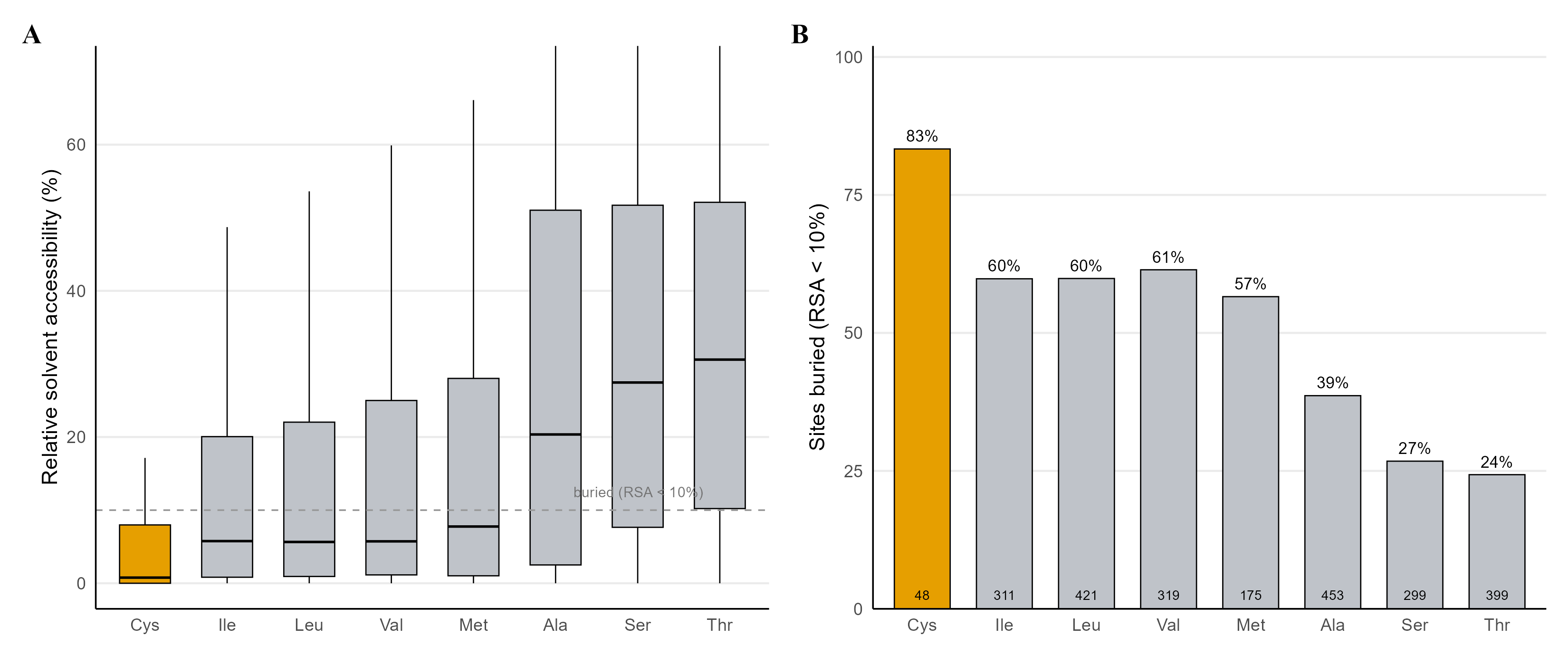
